# AAV-PHP.S-mediated delivery of reporters to cranial ganglion sensory neurons

**DOI:** 10.1101/2021.09.14.460327

**Authors:** Andoni I. Asencor, Gennady Dvoryanchikov, Pantelis Tsoulfas, Nirupa Chaudhari

**Affiliations:** Department of Physiology and Biophysics, University of Miami Miller School of Medicine, Miami, FL 33136, United States; Miami Project to Cure Paralysis, University of Miami Miller School of Medicine, Miami, FL 33136, United States; Department of Otolaryngology, University of Miami Miller School of Medicine, Miami, FL 33136, United States

**Author notes:** Correspondence to Nirupa Chaudhari. Author Contributions: AA, GD, PT and NC designed experiments, analyzed data and drafted/wrote the manuscript; AA, GD and PT performed the research; PT contributed unpublished reagents.

## Abstract

Because of their ease of use and low risk containment, Adeno-Associated Virus vectors are indispensable tools for much of neuroscience. Yet AAVs have been used relatively little to study the identities and connectivity of peripheral sensory neurons because methods to selectively target particular receptive fields or neuron types have been limited. The introduction of the AAV-PHP.S capsid with selective tropism for peripheral neurons (Chan et al., 2017) offered a solution, which we further elaborate here. We demonstrate using AAV-PHP.S with GFP or mScarlet reporters, that all cranial sensory ganglia, i.e. for cranial nerves V, VII, IX and X, are targeted. Pseudounipolar neurons of both somatic and visceral origin, but not satellite glia, are efficiently transduced rapidly and express the gene of interest within 1 week of injection. Fluorescent reporter proteins are transported into the central and peripheral axons of these sensory neurons, permitting visualization of terminals at high resolution, and/or in intact, cleared brain using light sheet microscopy. By combining a Cre-dependent reporter with the AAV-PHP.S capsid, we confirmed expression in a cell-type dependent manner for both anatomical and targeted functional analyses. The AAV-PHP.S capsid will be a powerful tool for mapping the receptive fields and circuits of molecular subtypes of many somatosensory, gustatory and visceral sensory neurons.

**SIGNIFICANCE STATEMENT:** AAV vectors have become an essential tool for visualizing, manipulating, and recoding from neurons of the central nervous system. However, the technology is not widely used for peripheral neurons because of several technical limitations. The AAV-PHP.S synthetic capsid, which targets peripheral neurons, was recently introduced (Chan et al., 2017). Here, we establish key parameters for using this virus, including which cells are transduced, the timing of expression in central and peripheral terminals, distant from neuronal somata, and the effectiveness of Cre-dependent constructs for cell type selective expression. This permits the use of AAV for constructing detailed anatomic and functional maps of the projections of molecular subtypes of peripheral sensory neurons.

## INTRODUCTION

Adeno-associated viruses (AAV), have emerged as one of the preferred tools of neuroscience research. Through their ease of use and effective delivery of various gene products including fluorescent proteins, AAVs have dramatically facilitated the mapping and manipulation of neural circuits in the brain (Samaranch et al., 2012; Nectow and Nestler, 2020). Depending on the serotype, AAVs are effective in different brain regions, display specific uptake into neuronal and/or glial populations, and are selectively transported in anterograde or retrograde direction in CNS neurons (Cearley et al., 2008; Ortinski et al., 2010; Salegio et al., 2013). Stereotaxic injections at target sites combined with the use of virally delivered or transgenically expressed Cre- and Flp-recombinases allows constructing detailed maps of the projections of selected neuronal populations and their functional interactions (Rothermel et al., 2013; Saunders and Sabatini, 2015; Weinholtz and Castle, 2021).

To date, AAV-dependent manipulation of the peripheral nervous system has been much more limited, although many open questions about connectivity would benefit from this approach. One difficulty is that the somata of such neurons are contained in relatively inaccessible dorsal root, cranial or autonomic ganglia. The peripheral terminals of sensory neurons are distributed in skin, muscle or viscera, making it difficult to target a defined anatomic or functional class of neurons. AAV-mediated transduction of peripheral neurons has been reported in a number of studies using various delivery methods such as sciatic nerve injection (Towne et al., 2009), direct intraganglionic injection (Kollarik et al., 2010) and intrathecal infusion (Vulchanova et al., 2010; Schuster et al., 2013). However, these approaches are invasive, and inefficient the viral particles lack wide tropism to peripheral neurons. Another approach for preferentially transducing peripheral sensory neurons has been to inject virus into the circulation, taking advantage of the poor transport of AAVs across the blood-brain barrier (Watson et al., 2016; Bloom et al., 2019). By itself, this did not fully limit transduction to peripheral neurons. A major advance came with the development of the synthetic neurotropic serotype, AAV.PHP.S, which was produced by a directed evolution. Variations in the Cap gene of AAV9 were introduced, followed by iterative phenotypic selection for transduction of dorsal root ganglion sensory neurons (Chan et al., 2017). The resulting pseudotyped AAV-PHP.S particles, when introduced into the circulation, infected mostly or only peripherally located neurons.

Although Chan et al reported that the AAV-PHP.S capsid displays tropism towards dorsal root ganglion and enteric neurons, they did not elaborate on other cranial ganglia with their distinct sensory neuron types. Nor were the projections of these peripheral sensory neurons examined in the original report. Here, we have systematically explored and report on transduction of neurons in the trigeminal, geniculate, petrosal and nodose ganglia (cranial nerves V, VII, IX and X) using AAV-PHP.S. We also report on the time course for the expression of fluorescent reporters in neuronal somata and the transport of fluorescence to peripheral and central terminals, which are at considerable distance. Finally, we demonstrate using GCaMP and both functional and anatomical criteria, that PHP.S viral particles can deliver reporters for stringent Cre-dependent expression across a range of viral titers, permitting exhaustive or sparse labeling according to experimental needs.

## METHODS

### Animals and Tissues

All experiments were carried out according to the NIH Guidelines for the Care and Use of Laboratory Animals, and protocols were approved by the University of Miami Institutional Animal Care & Use Committee. Mice of the following strains (stock #) were purchased from The Jackson Laboratory and bred in-house: C57BL/6J (#000664), *Mafb*-2A-mCherry-2A-Cre (#029664) (Wu et al., 2016). Additionally, *Pirt*-Cre knockin mice, in which Cre is expressed in all sensory ganglion neurons (Kim et al., 2008) were a kind gift from X. Dong, Johns Hopkins University, and were crossed with RCL-GCaMP6s (Jax Ai96, #024106) to produce *Pirt*;;GCaMP6s. Mice of the *Plcb2*-GFP strain (Kim et al., 2006) were produced and bred in-house.

### Plasmids and AAV

Plasmids for producing AAV particles for GFP or GCaMP expression were obtained from Addgene (Watertown, MA), including pAAV-CAG-GFP (#37825), a gift from Edward Boyden, and pAAV.CAG.Flex.GCaMP6s.WPRE.SV40 (#100842), a gift from Douglas Kim. The cDNA for the red fluorescent protein, mScarlet-I (NCBI #KY021424) (Bindels et al., 2017), was optimized based on human codon usage, then synthesized by GeneArt (ThermoFisher). This cDNA was used to replace the GFP sequence in pAAV-CAG-GFP (above) at BamHI and EcoRV sites.

Viruses were produced at the University of Miami viral core facility at the Miami Project to Cure Paralysis, using pUCmini-iCAP-PHP.S (Addgene #103006) in HEK293T cells. Titers of AAV-PHP.S preparations (in viral genome copies / ml, assessed by qPCR) were as follows: CAG-GFP, 1.3×10^14^; CAG-mScarlet-I, 3.8×10^14^; and CAG-Flex-GCaMP6s, 2.9×10^14^.

Male or female mice between 2 and 6 months of age were retro-orbitally injected with ≈1.3 to 3.8×10^12^ viral particles per mouse, using Terumo 1cc tuberculin syringes with 26G × 3/8” needle. For the time course and titer series, mice of both sexes were randomly assigned.

### Immunohistochemistry and imaging

For perfusion-fixation, mice were deeply anesthetized with ketamine and transcardially perfused sequentially with cold saline (0.9% NaCl), then 4% paraformaldehyde in saline. Tissues were dissected, post-fixed for 1 hour at 4°C (except overnight for brain and spinal cord), washed in phosphate-buffered saline (PBS: 3.8 mM NaH_2_PO_4_, 16.2 mM Na_2_HPO_4_, 15 mM NaCl/ 1L), cryoprotected overnight at 4°C in 30% sucrose in PBS, and embedded in OCT.

Cryosections were cut on a Leica CM1900, sensory ganglia and lingual tissue to 20 µm thickness, spinal cord and brain to 40 µm. Sections, mounted on slides, were permeabilized (0.1% Triton X-100 in PBS), blocked (10% normal donkey serum) and incubated in diluted primary antibodies overnight at 4°C. After washing for an hour in PBS, secondary antibody was incubated for 1-2 h. Sections were mounted under Fluoromount G (SouthernBiotech, Birmingham, AL). Antibodies used and their concentrations are in **Table 1**.

**Table 1.**
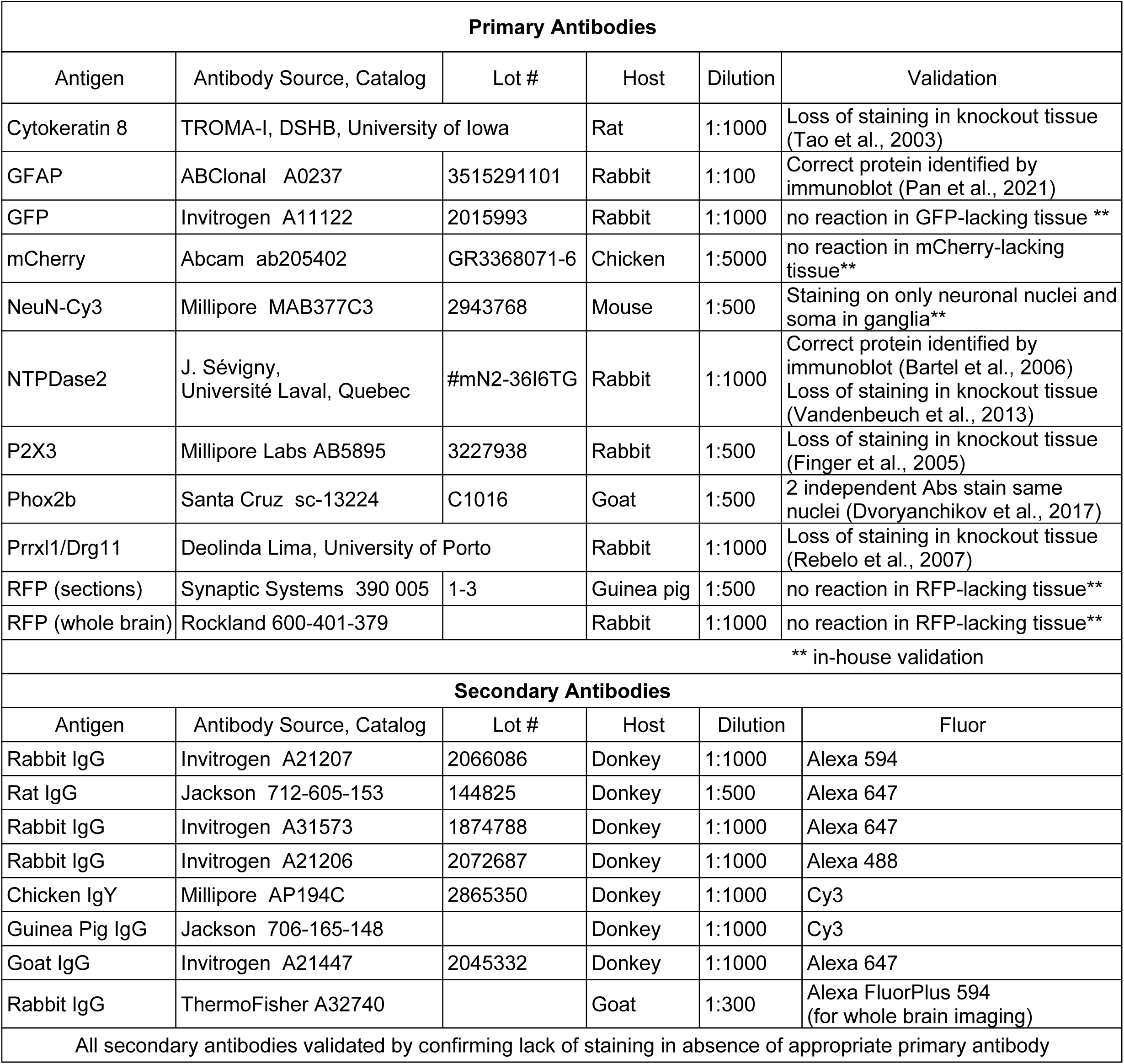
Primary and secondary antibodies used.

| Primary Antibodies |  |  |  |  |  |
| --- | --- | --- | --- | --- | --- |
| Antigen | Antibody Source, Catalog | Lot # | Host | Dilution | Validation |
| Cytokeratin 8 | TROMA-I, DSHB, University of Iowa |  | Rat | 1:1000 | Loss of staining in knockout tissue (Tao et al., 2003) |
| GFAP | ABClonal A0237 | 3515291101 | Rabbit | 1:100 | Correct protein identified by immunoblot (Pan et al., 2021) |
| GFP | Invitrogen A11122 | 2015993 | Rabbit | 1:1000 | no reaction in GFP-lacking tissue ** |
| mCherry | Abcam ab205402 | GR3368071-6 | Chicken | 1:5000 | no reaction in mCherry-lacking tissue** |
| NeuN-Cy3 | Millipore MAB377C3 | 2943768 | Mouse | 1:500 | Staining on only neuronal nuclei and soma in ganglia** |
| NTPDase2 | J. Sévigny, Université Laval, Quebec | #mN2-36I6TG | Rabbit | 1:1000 | Correct protein identified by immunoblot (Bartel et al., 2006)<br>Loss of staining in knockout tissue (Vandenbeuch et al., 2013) |
| P2X3 | Millipore Labs AB5895 | 3227938 | Rabbit | 1:500 | Loss of staining in knockout tissue (Finger et al., 2005) |
| Phox2b | Santa Cruz sc-13224 | C1016 | Goat | 1:500 | 2 independent Abs stain same nuclei (Dvoryanchikov et al., 2017) |
| Prrxl1/Drg11 | Deolinda Lima, University of Porto |  | Rabbit | 1:1000 | Loss of staining in knockout tissue (Rebelo et al., 2007) |
| RFP (sections) | Synaptic Systems 390 005 | 1-3 | Guinea pig | 1:500 | no reaction in RFP-lacking tissue** |
| RFP (whole brain) | Rockland 600-401-379 |  | Rabbit | 1:1000 | no reaction in RFP-lacking tissue** |
| ** in-house validation |  |  |  |  |  |
| Secondary Antibodies |  |  |  |  |  |
| Antigen | Antibody Source, Catalog | Lot # | Host | Dilution | Fluor |
| Rabbit IgG | Invitrogen A21207 | 2066086 | Donkey | 1:1000 | Alexa 594 |
| Rat IgG | Jackson 712-605-153 | 144825 | Donkey | 1:500 | Alexa 647 |
| Rabbit IgG | Invitrogen A31573 | 1874788 | Donkey | 1:1000 | Alexa 647 |
| Rabbit IgG | Invitrogen A21206 | 2072687 | Donkey | 1:1000 | Alexa 488 |
| Chicken IgY | Millipore AP194C | 2865350 | Donkey | 1:1000 | Cy3 |
| Guinea Pig IgG | Jackson 706-165-148 |  | Donkey | 1:1000 | Cy3 |
| Goat IgG | Invitrogen A21447 | 2045332 | Donkey | 1:1000 | Alexa 647 |
| Rabbit IgG | ThermoFisher A32740 |  | Goat | 1:300 | Alexa FluorPlus 594<br>(for whole brain imaging) |
| All secondary antibodies validated by confirming lack of staining in absence of appropriate primary antibody |  |  |  |  |  |

Imaging was on an Olympus Fv1000 BX61 upright or an Olympus Fv1000 IX81 inverted laser scanning confocal microscope. Multi-channel images were captured and were adjusted, only for brightness, in Photoshop. No contrast enhancement was applied. All images are shown at, or smaller than captured size. For the 3-D imaging, we used a Miltenyi Biotec UltraMicroscope (see below).

### Cell Counting

To quantify the efficiency of delivery of GFP, 20μm thick cryosections of geniculate ganglia were evaluated by intrinsic fluorescence of GFP and immunofluorescence for NeuN. A stack of confocal images was captured for each tissue section. To ensure that each GFP+ neuron was counted only once, we set background fluorescence as the intensity on the adjacent facial nerve (motor) track, viewed neurons in Z-projection so that neuronal shape and size (15-25μm diameter) was evident and included a NeuN-immunoreactive nucleus. With these criteria, neurons which displayed faint to bright fluorescence were considered GFP+; many other NeuN+ neurons were clearly GFP-negative.

### *In vivo* Ca^2+^ imaging

Mice were anesthetized with ketamine and xylazine, the geniculate ganglion was exposed via a dorsal approach and imaged for GCaMP6s fluorescence as previously described (Wu et al., 2015; Leijon et al., 2019). Mechanical stimuli to the pinna included puffs of compressed air or stroking with a bristle brush for 10s. Taste stimuli were perfused through the oral cavity as described (Wu et al., 2015; Leijon et al., 2019), and included 300mM sucrose (sweet), 250mM NaCl (salty), 10mM citric acid (sour), a mix of 1µm cycloheximide + 0.3mM quinine (bitter), 20mM monosodium glutamate + 1mM inosine monophosphate (umami) in sequence. Time series of fluorescence images were analyzed using ImageJ and are presented as changes of fluorescence normalized to baseline (*i*.*e*., ΔF/F), for individual Regions of Interest, each representing a single neuron.

### Tissue Clearing and Light Sheet Microscopy

For whole brain staining and clearing, we used an enhanced version of iDISCO (Renier et al., 2014; Bray et al., 2017; http://lab.rockefeller.edu/tessier-lavigne/assets/file/whole-mount-staining-bench-protocol-january-2015.pdf). Dissected whole brain was dehydrated through a methanol/PBS series, bleached overnight at 4°C, rehydrated, permeabilized at 37°C for 2 d, blocked for 2 d and then incubated with anti-RFP at 37°C for 10 days. After washing overnight, the brain was incubated in secondary antibody for 10 days, then washed, dehydrated and cleared as described previously.

After clearing, samples were imaged the same day using light-sheet microscopy (Ultramicroscope, LaVision BioTec) using a fluorescence macro zoom Olympus MVX10 microscope with a 2× Plan Apochromatic zoom objective (NA 0.50). Image analysis and 3D reconstructions were performed using Imaris v9.5 software (Bitplane, Oxford Instruments) after removing autofluorescence using the Imaris Background Subtraction function with the default filter width so that only broad intensity variations were eliminated.

## RESULTS

### Selectively targeting sensory ganglion neurons

Gene delivery to dorsal root ganglion neurons was previously demonstrated using the PHP.S synthetic capsid. To explore whether this capsid effectively targets both somatic and visceral sensory ganglion neurons, we injected the equivalent of ≈10^12^ viral genomes of AAV.PHP.S::CAG-GFP into the retroorbital sinus of mice and after 1 week, examined the trigeminal (cranial V), geniculate (cranial VII), petrosal (cranial IX), nodose (cranial X), and several dorsal root ganglia. The trigeminal and dorsal root ganglia confer general somatic sensation in the head and neck and the rest of body respectively while the petrosal and nodose are visceral sensory ganglia that innervate tastebuds of the posterior tongue and pharynx, and multiple viscera. The geniculate ganglion is hybrid, including both general somatic sensitivity for the pinna and a visceral contingent of gustatory neurons innervating tastebuds of the anterior tongue (D’Autreaux et al., 2011; Dvoryanchikov et al., 2017).

In cryosections of all the above ganglia, 7 days after injection and later, we observed substantial expression of GFP. However, the density of GFP+ cells varied, even when different ganglia from a single mouse were directly compared (**Fig. 1A-D**). The brightness of GFP fluorescence across neurons within a single ganglion was also variable, which may reflect different numbers of viral particles infecting individual cells, or that some neuron types express the reporter more efficiently or rapidly.

**Figure 1.**
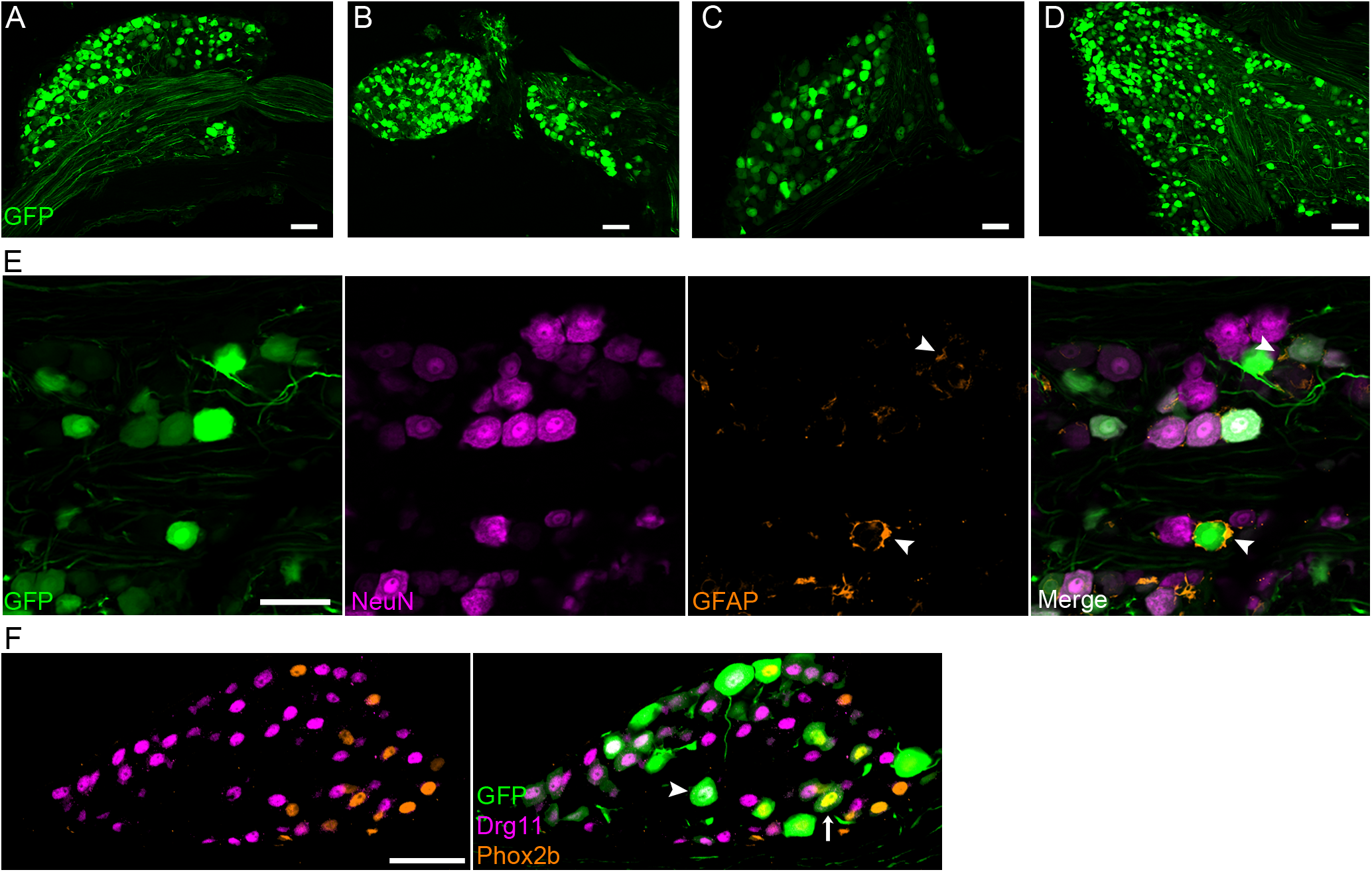
AAV-PHP.S transduces neurons in multiple sensory ganglia. AAV-PHP.S::CAG-GFP, injected into the retroorbital sinus, resulted in expression of GFP in **A**, dorsal root, **B**, nodose-petrosal complex, **C**, geniculate, and **D**, trigeminal ganglia, viewed for GFP intrinsic fluorescence in cryosections. Ganglia were dissected 7 days post-injection. **E**, Cryosections of trigeminal ganglion (as in D) were immunostained for NeuN (magenta) and GFAP (orange) to identify neurons and satellite glia respectively. Only neurons are seen to express GFP. **F**, Cryosections of geniculate ganglion (as in **C**) were immunostained for Phox2b (orange) and Drg11 (magenta) to identify visceral and somatic neurons respectively. Neurons of both classes (arrow, arrowhead) express GFP (green). All images are confocal single planes. Scale bars, 50μm.

To evaluate if AAV.PHP.S was targeting glia in addition to neurons, we co-immunostained sections of trigeminal ganglion for NeuN to detect neurons and GFAP, expressed at low levels in a subset of satellite glia which encase the ganglion neurons (Chudler et al., 1997; Villa et al., 2010). The distinctive crescent shape of satellite glia was not observed among GFP+ cells. GFP-expressing cells were consistently NeuN+, large and polygonal, and GFAP-negative (**Fig. 1E**). We also confirmed that GFP-expressing cells in the geniculate, petrosal, nodose and dorsal root ganglia also were consistently shaped and sized like neurons and NeuN+ (not shown). Thus, AAV-PHP.S appears to target neurons of a broad range of distinct sensory ganglia, but not their satellite cells.

To assess if both somatic and visceral sensory neurons were transduced, we immunostained geniculate ganglion cryosections for Phox2b and Drg11/Prrxl1, respectively markers for the gustatory visceral neurons and for the somatosensory auricular neurons in this ganglion (D’Autreaux et al., 2011; Dvoryanchikov et al., 2017). Both classes were found among the GFP+ transduced neurons (arrows and arrowheads in **Fig.1F**).

We did not observe GFP-label in spiral ganglion neurons of the cochlea, nor in the olfactory epithelium (not shown).

### Maximal expression in ganglia occurs by 7 days

To explore the time-course for GFP expression by AAV-PHP.S, we selected the geniculate ganglion, as it contains both somatic and visceral sensory neurons (D’Autreaux et al., 2011). We retro-orbitally injected 8 mice, each with 10^12^ viral genome, and examined them across a 3-week time-course (**Fig. 2A-E**). As early as 2 days post-injection, traces of GFP fluorescence could be detected in a few geniculate ganglion neurons. To quantify the efficiency of reporter delivery, we immunostained all cryosections of individual geniculate ganglia for NeuN and scored the fraction of NeuN+ cells that were GFP-labeled. By 7 days post-injection and subsequently, the fraction visible as GFP+ stabilized at ≈65% of all neurons in the ganglion. The peak fraction of expressing cells at 7 days for PHP.S is similar to time frame reported for other strains of AAV (Zincarelli et al., 2008).

**Figure 2:**
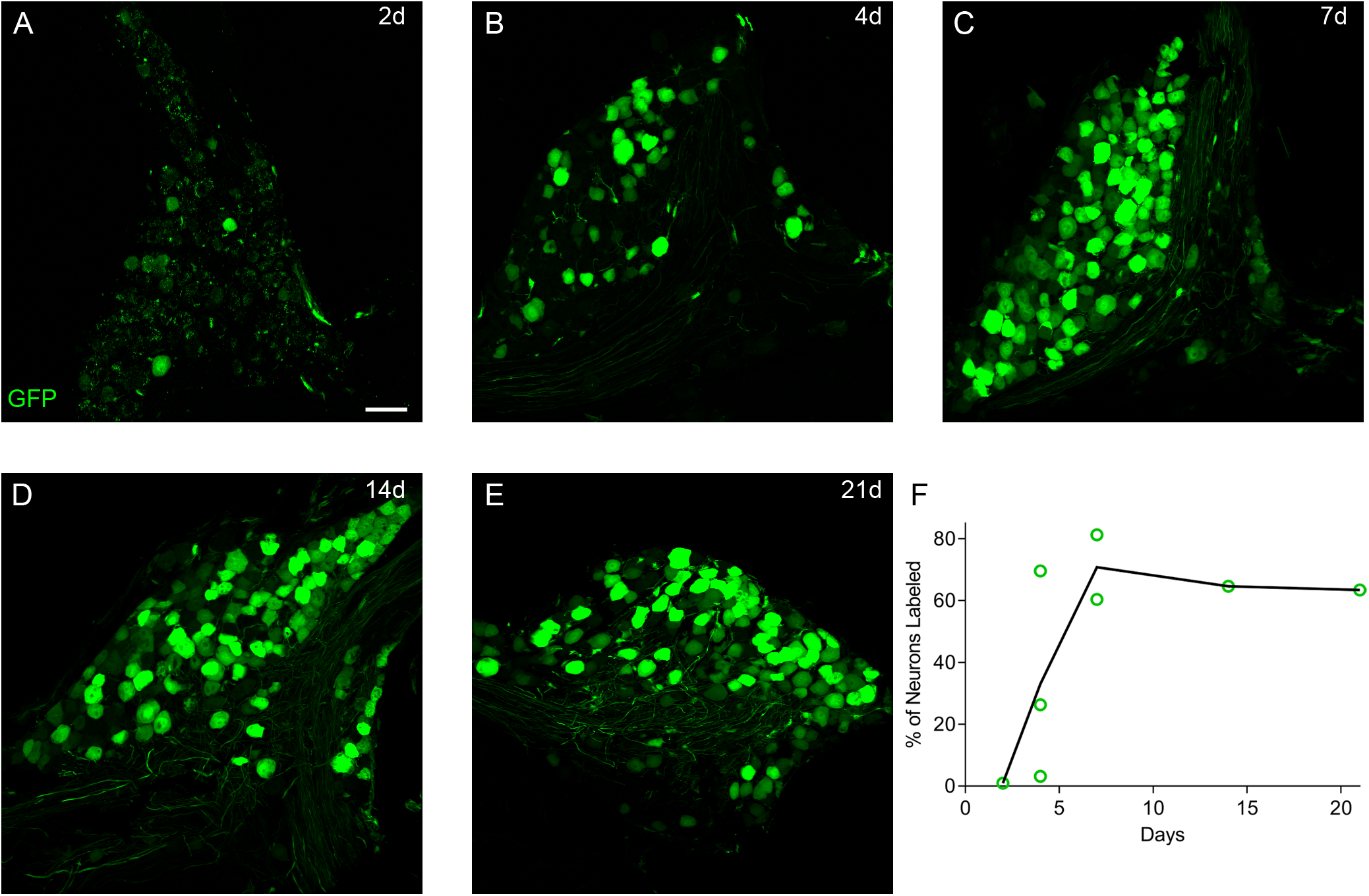
Time-course of GFP expression. **A-E**, Cryosections of geniculate ganglia from mice injected with AAV-PHP.S::CAG-GFP, and analyzed at 2, 4, 7, 14, and 21 days post-injection. Intrinsic fluorescence of GFP was captured in parallel for all images; brightness was increased in **A** to visualize the very low level expression at the earliest stage. **F**, transduction efficiency was quantified in confocal images of cryosections stained for NeuN, as the fraction of NeuN+ cells that were GFP+. Symbols represent individual mice; solid line is average at each time point. All images are confocal, Z-projected over 10-14μm. Scale bar, 50μm.

### CNS neurons remain unlabeled

The use of the PHP.S capsid along with injection into the general circulation was reported to limit viral targeting to the peripheral nervous system (Chan et al., 2017). However, the report did not elaborate on how effective this restriction was. If this virus is to be useful for tracing the central projections of sensory afferent neurons, it is essential that there should be little to no labeling of resident neurons in areas where such central projections terminate. Thus, we examined the brainstem and spinal cord in areas that contain the central projections of sensory neurons targeted by AAV-PHP.S.

In hindbrain sections, GFP labeled axons were clearly detected in spinal trigeminal tract (Sp5) and in the Nucleus of the Solitary Tract (NST) (**Fig. 3A**) which contain respectively, the central projections of sensory neurons from the trigeminal ganglion or the ganglia of the VII, IX, X cranial nerves (Hamilton and Norgren, 1984). In the spinal cord also, GFP label was detected in the dorsal horn, with additional GFP+ fibers extending ventrally past the central canal (**Fig.3B**). At higher magnification, both Sp5 and NST contained GFP+ fibers with occasional boutons that appear similar to synapses. Importantly, we found no resident neuronal somata that were GFP-labeled in either of these two regions (**Fig.3C**,**D**). In the rostral NST, where gustatory afferents project, many GFP+ fibers were immunoreactive for P2×3, a known marker for gustatory neurons (Bo et al., 1999; Finger et al., 2005).

**Figure 3:**
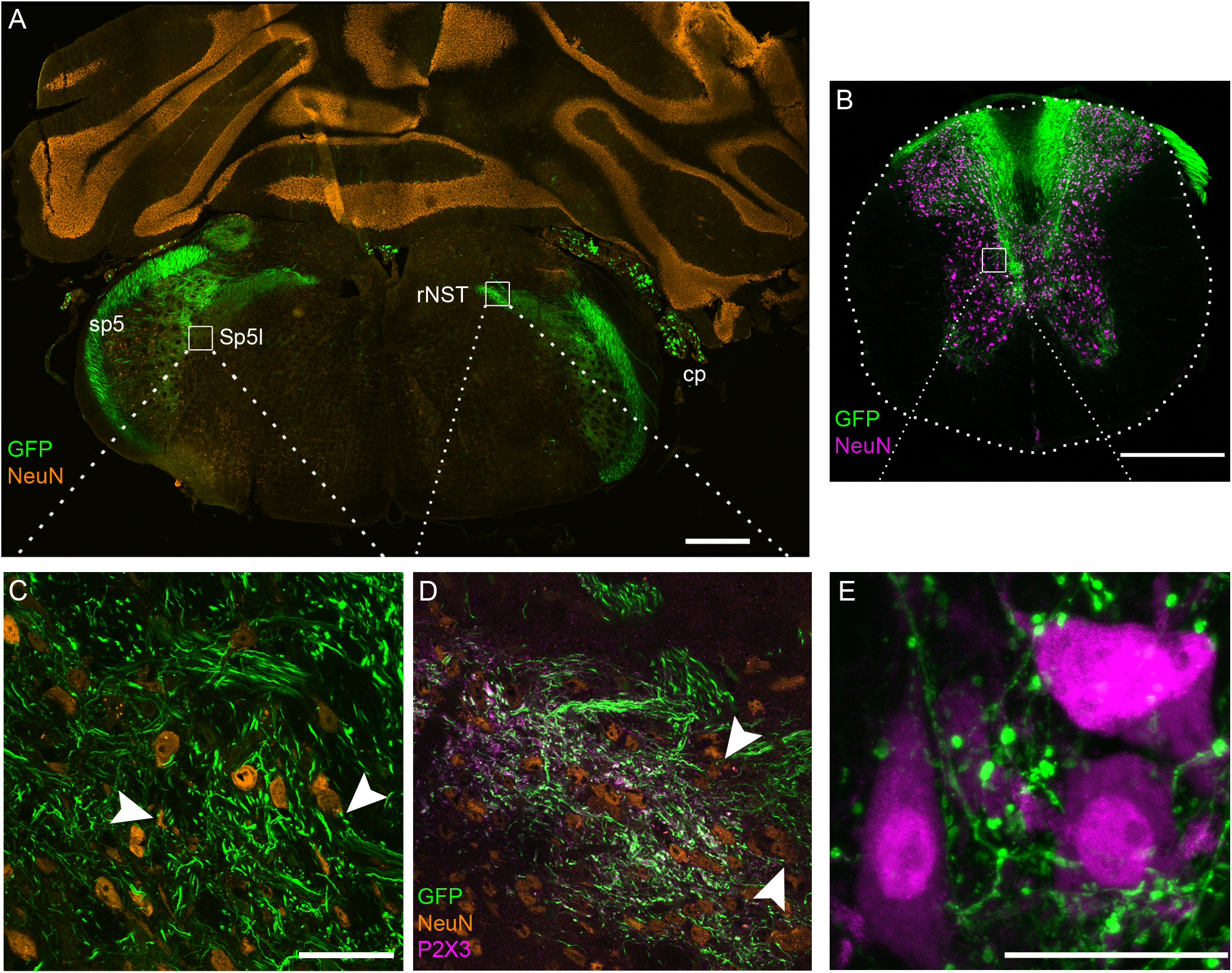
Only PNS neurons are targeted; CNS neurons remain unlabeled. Mice were injected with AAV-PHP.S::CAG-GFP and tissues were harvested 14 days later. **A**, Cryosection of hindbrain, immunostained for NeuN (orange) shows that AAV-delivered GFP is limited to the choroid plexus (cp) and areas that receive afferent inputs from peripheral sensory neurons: the spinal trigeminal tract (sp5), spinal trigeminal nucleus interpolar (Sp5l) and rostral Nucleus of Solitary Tract (rNST) with its gustatory inputs. **B**, section of mid-thoracic spinal cord, imaged for NeuN immunoreactivity (magenta) and intrinsic fluorescence of AAV-delivered GFP, which is limited to sensory inputs from the dorsal root ganglia. **C**, single confocal plane, higher magnification of boxed region of Sp5l showing the lack of GFP+ central neurons, and fine GFP-labeled sensory fibers, some of which terminate on neurons in this region of the trigeminal nucleus (arrowheads). **D**, single confocal plane at higher magnification of boxed region of rNST, additionally stained for P2×3 (magenta) to label gustatory afferents. Many of the finest GFP+ fibers are gustatory and appear to synapse on rNST neurons (arrowheads). **E**, high magnification of **B** in ventral horn boxed region where GFP+ afferent sensory fibers (proprioceptors) synapse onto primary motor neurons. Scale bars, A, B: 500μm; C-E: 50μm.

In the spinal cord, as well, there were no GFP+ neuronal somata in the grey matter. Instead, GFP+ fibers from the dorsal root ganglia terminated throughout the dorsal horn. A minority of GFP+ fibers continued ventrally to terminate on very large neuronal somata (**Fig.3E**), representing proprioceptor fibers projecting to motor neurons in the ventral horn.

Qualitatively similar labeling of central terminals was detected at 7- and 14-days post-injection, although the GFP-labeled fibers appear to be more numerous and somewhat brighter at 14 days.

### Sensory afferent fibers distant from soma are labeled

Because high levels of GFP could be detected in ganglia by 7 days, we examined if the fluorescent reporter could be detected several mm away from the soma, in the sensory peripheral terminals. We selected for analysis, fungiform and circumvallate taste buds, which are located 1-2 cm from the somata of the neurons that innervate them. Cryosections of lingual papillae from mice injected with AAV.PHP.S:: CAG-GFP virus were immunostained for Ker8 and P2×3 to visualize taste buds and gustatory afferent fibers respectively. At 7 days post-injection, some nerve fibers in circumvallate taste buds exhibited faint fluorescence. In contrast, fungiform taste buds, which are as much as 1cm farther from the somata of their innervating neurons showed no GFP+ fibers within them (**Fig.4A**). Over the course of two additional weeks (**Fig.4B**,**C**), more fibers became GFP-labeled within the taste buds, and these displayed coexpression with P2×3. We noted that across all tissues and images, the fibers defined by expressed GFP reporter were finer and better resolved when compared to immunostained fibers.

**Figure 4:**
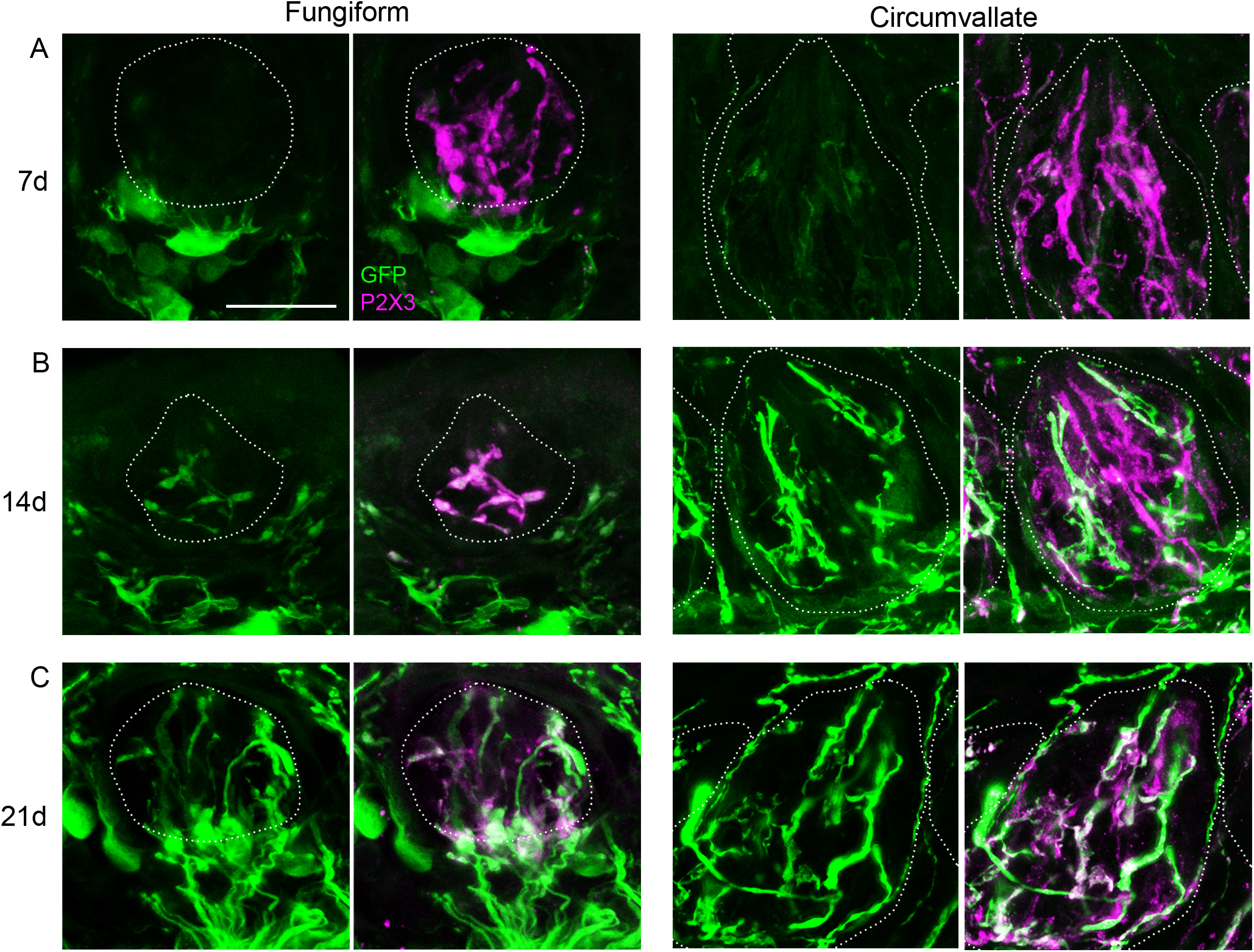
Peripheral terminals of sensory ganglion neurons are GFP-labeled. **A-C**. Tongues from mice, injected with AAV-PHP.S::CAG-GFP were examined at 7, 14 or 21 days post-injection to examine arrival of axonally transported GFP to the peripheral terminals of trigeminal, geniculate, and petrosal ganglion neurons. Dotted lines outline tastebuds. Gustatory fibers are immunoreactive for P2×3 (magenta) while trigeminal fibers, outside the taste buds,remain unstained. **A**, in fungiform taste buds at 7 days (left), GFP has not yet been transported into gustatory fibers (magenta), while in circumvallate taste buds, gustatory fibers show faint GFP fluorescence. **B**,**C**, both gustatory and trigeminal fibers show progressively brighter GFP labeling over time in both fungiform and circumvallate taste buds. All images were captured in parallel at the same settings to make fluorescence intensities comparable. Each panel is a Z-projection across ≈5μm thickness. Scale bar, 20μm.

In lingual sections (e.g. **Fig.4A**), we observed large GFP-expressing cells located in connective tissue below the taste bud and epithelium. Their location and morphology suggest resident macrophages or dendritic cells, although we did not characterize them further. These cells became GFP+ well before GFP is detected in fibers innervating the epithelium.

We also tested another fluorescent reporter, mScarlet-I, which is bright and highly suitable for imaging cleared whole brain, and offers a second color for dual-label transductions. We injected AAV.PHP.S:: CAG-mScarlet into *Plcb2-*GFP mice in which the type II chemosensory cells of taste buds express GFP (Kim et al., 2006). Interactions of individual type II cells with individual afferent fibers could readily be visualized in fungiform (**Fig. 5A**) and circumvallate (**Fig.5B**,**C**) taste buds. In both locations, boutons on afferent fibers as they terminate on the type II cells are readily seen (arrow). If either AAV-delivered GFP or mScarlet-I were expressed selectively in a single neuron type from the ganglion, the method would permit precise definition of neuron-target interactions.

**Figure 5:**
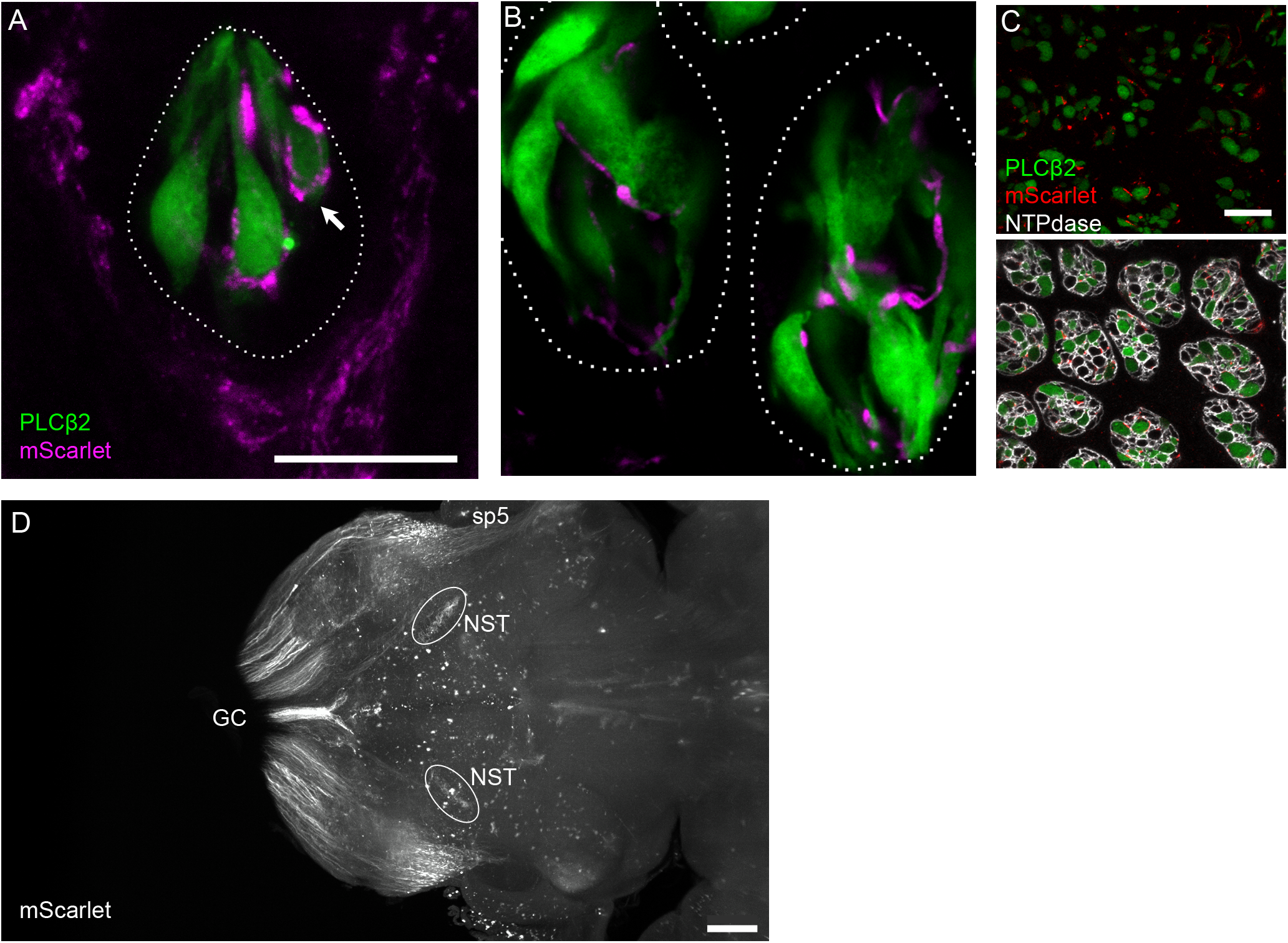
mScarlet-I reporter for visualizing individual fibers and fiber tracts. **A-C**, fungiform and circumvallate tastebuds from a *Plcb2*-GFP mouse, injected with AAV-PHP.S::CAG-mScarlet, examined 21 days post-injection. In the fungiform papilla (**A**), two types of mScarlet+ nerves (magenta) are visible: trigeminal fibers that form a corona around the taste bud, and gustatory fibers that enter the taste bud. Terminal boutons of gustatory nerves are seen associated with individual type II (GFP-expressing, green) chemoreceptor cells (arrow). **B**, In a single confocal plane, two circumvallate taste buds similarly show gustatory afferents associating with GFP+ chemosensory cells. **C**, circumvallate taste buds from the same mouse as in A and B, viewed in cross-section and at low magnification, Nearly every taste bud includes one or more afferent fibers that are mScarlet-labeled (red). Immunoreactivity for NTPDase2 outlines each taste bud cell. **D**, Light-sheet microscope imaging of a cleared brain of a mouse injected with AAV-PHP.S;;CAG-mScarlet. Brain was viewed in a horizontal orientation. This image is a Z-projection through a 100 μm thickness that includes the rostral Nucleus of the Solitary Tract (NST). Sensory afferent fibers of AAV-transduced neurons detected here include the gracile and cuneate sensory tracts (GC) entering the medulla caudally, the spinal trigeminal tract (sp5) and the terminals of gustatory afferents in the rostal nucleus of the solitary tract (NST). Scale bar, 20μm for taste buds, 500μm for brain.

To visualize primary sensory fiber tracts in the brain in a 3-dimensional manner, we subjected intact brains from mice injected with AAV.PHP.S:: CAG-mScarlet-I to immunostaining to enhance mScarlet fluorescence, followed by clearing and imaging by light sheet microscopy (**Fig. 5D**). Ascending spinal sensory tracts as well as trigeminal and gustatory tracts could readily be traced from their entrance into the CNS to their terminals in respective sensory nuclei (**Fig. 5D** and **Extended Data-1**).

### Cell type selective expression and function

To selectively target subtypes of peripheral sensory neurons in the geniculate ganglion, we employed a Cre-dependent construct, AAV-PHP.S::CAG-flex-GCaMP6s and injected ≈10^12^ viral genomes into each of 2 *Mafb-*mCherry-Cre knockin mice. In the geniculate ganglion of this strain, Mafb is expressed at high levels in T2 gustatory neurons and at low levels in auricular neurons (Dvoryanchikov et al., 2017). Accordingly, these two neuron types are distinguishable by bright or faint red intrinsic mCherry fluorescence respectively. The large majority of gustatory neurons are mCherry-negative. Cryosections were immunostained with an antibody against Phox2b, to identify gustatory neurons (Dvoryanchikov et al., 2017). Approximately half of all T2 and auricular neurons were found to express GCaMP (white arrowheads and arrows, respectively, **Fig. 6**). In the same ganglia, fewer than 10% of neurons which lack Mafb, mCherry, and Cre expressed GCaMP. Similar results regarding incidence and specificity of transduced neurons were obtained at 10 and 40 days post-injection. Cre-dependence was also confirmed by the lack of GCaMP expression in the geniculate ganglion of a AAV-PHP.S::CAG-flex-GCaMP6s-injected mouse that lacked the Cre transgene (data not shown).

**Figure 6:**
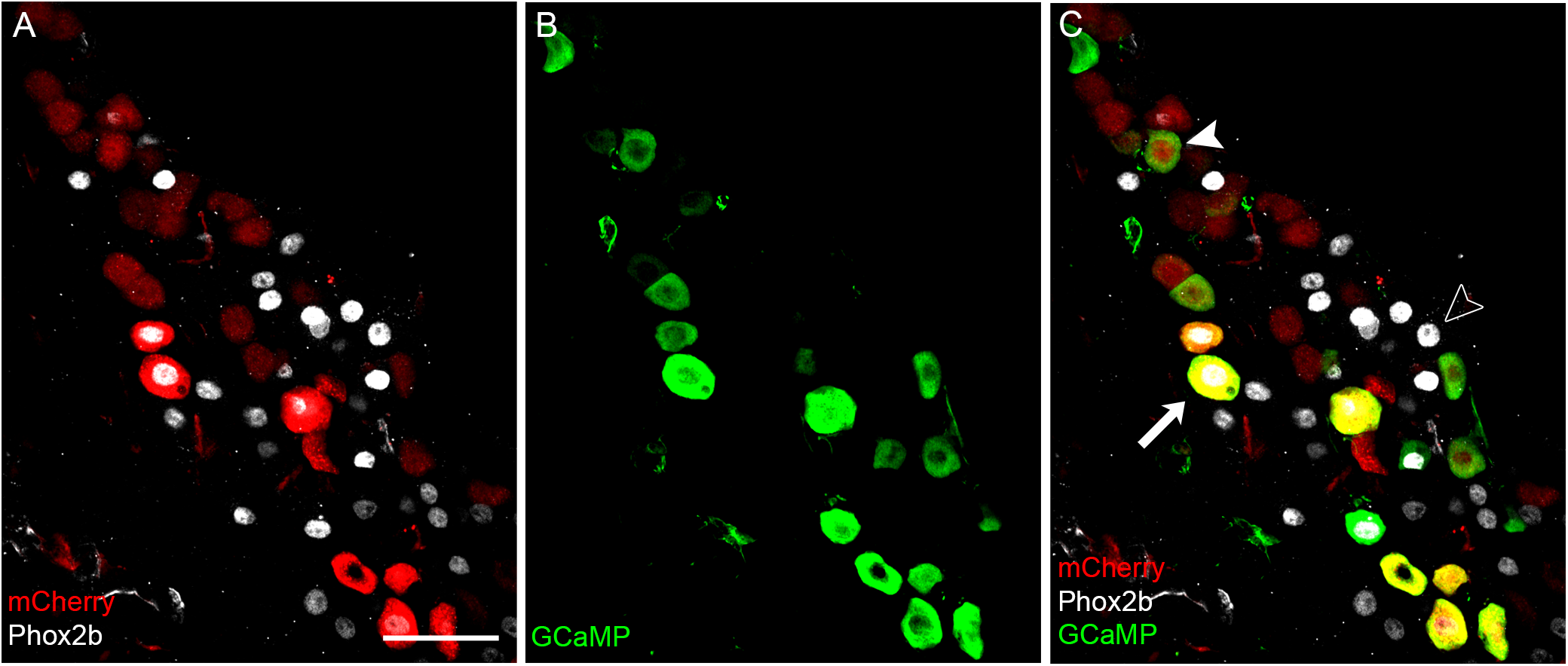
Cre-dependent expression in peripheral sensory neurons. Geniculate ganglion of a *Mafb*-mCherry-Cre mouse (in which Cre and mCherry are co-expressed) was injected with AAV-PHP.S::*flex*-GCaMP6s and examined 10 days post-injection. Cryosections were immunostained with anti-GFP (to detect GCaMP), anti-mCherry, and anti-Phox2b (to detect gustatory neurons). GCaMP6 (green) is detected predominantly in neurons that express Cre and mCherry (red). The ganglion includes two classes of Cre-expressing neurons: faint red, Phox2b-negative auricular neurons and large, bright red, Phox2b+ T2 gustatory neurons (Dvoryanchikov et al., 2017). Examples of these two types, expressing in Cre-dependent manner are indicated by a white arrowhead or arrow respectively. Cre-lacking, Phox2b+ gustatory neurons do not express GCaMP (open arrowhead). Scale bar, 50um.

We tested if multiplicity of infection influences the Cre-dependence of AAV-PHP.S::CAG-Flex-GCaMP6s by injecting 4 *Mafb-*mCherry-Cre mice with a whole body load of 30, 15, 7.5 or 3.8×10^11^ genomes per animal. Geniculate ganglia from these mice were processed as in Fig. 6. At the lowest titer, no GCaMP expression was detected in any neurons. However, at all three of the higher titers, GCaMP expression was overwhelmingly (>90%) in mCherry+ (i.e. Cre-expressing) neurons (data not shown).

We verified that AAV-PHP.S::CAG-Flex-GCaMP6s targeted only peripheral neurons by examining hindbrain and spinal cord of *Mafb-*mCherry-Cre mice injected with the virus. Although a subset of neurons in the ventral cochlear nucleus and spinal trigeminal nucleus of the hindbrain and in the dorsal horn of the spinal cord do express Mafb (Lein et al., 2007; http://mouse.brain-map.org/gene/show/16430) such central neurons remained unlabeled with GCaMP6 (**Extended Data Fig. 2**).

Finally, we also confirmed that GCaMP, delivered by AAV-PHP.S to peripheral sensory neurons of *Mafb*-mCherry-Cre mice can be used for functional imaging. Using *in vivo* Ca^2+^ imaging as we previously described (Wu et al., 2015; Leijon et al., 2019), we recorded the responses of geniculate ganglion neurons (**Fig. 7A**) to gustatory and auricular stimuli including brushing the pinna with stiff bristles and washing a mix of tastants through the oral cavity. Auricular neurons in injected mice responded robustly and repeatedly to brushing the pinna, but not to tastants in the mouth (**Fig. 7B)**. Importantly, no neurons responded to taste stimuli, confirming the specificity of GCaMP targeting to only Mafb+ (i.e. non-taste) neurons.

**Figure 7:**
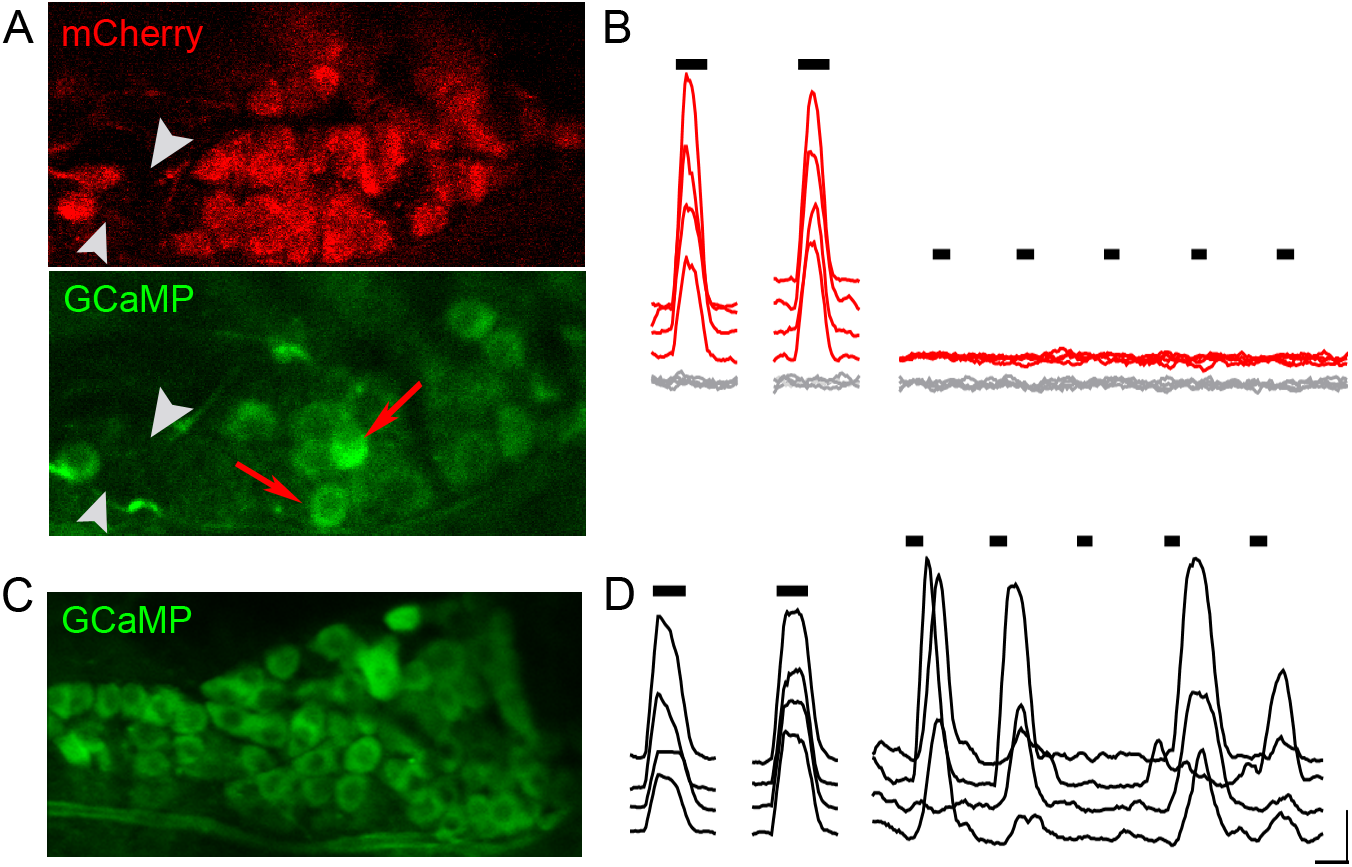
*In vivo* Ca^2+^ imaging with PHP.S delivered Cre-dependent GCaMP. **A**. Geniculate ganglion as in Fig.6, of anesthetized mouse, 19 days post-injection. Viewed for mCherry (red) or GCaMP (green) during recording. This region includes a high density of auricular neurons. **B**. Red traces are the responses (ΔF/F_0_) of 4 mCherry+ auricular neurons (2 examples identified by red arrows in **A**) to stimuli (black bars) from left to right: pinna stimulated with a puff of compressed air, a bristle brush, followed by oral perfusion of five tastants. Areas of the ganglion that contained mCherry-neg neurons (2 examples identified by grey arrowheads in **A**) showed no changes in baseline fluorescence during the same stimulation interval (grey traces). **C**. Geniculate ganglion from a *Pirt*;;GCaMP6s mouse in which all neurons are GCaMP+. **D**, Traces from ganglion in **C**, from 4 representative neurons each of auricular (left) and gustatory (right) classes stimulated in similar fashion as in **B**. Scale bars for B and C, 20s, 1.0 ΔF/F_0_.

In contrast, the same auricular and taste stimuli, delivered to a *Pirt*;;GCaMP6s mouse (GCaMP expressed transgenically in all geniculate ganglion neurons, not by viral transduction), elicited robust responses from appropriate auricular and gustatory neurons as predicted (shown for comparison in **Fig. 7C**).

## DISCUSSION

The creation of the AAV-PHP.S capsid (Chan et al., 2017) introduced the possibility of transducing peripheral neurons with a variety of reporters and/or gene products for anatomical and physiological studies. We have elaborated on the original study by demonstrating that in addition to dorsal room ganglion and enteric neurons, the neurons of many cranial ganglia, including both visceral and somatic classes also can be transduced. We show that virally delivered GFP becomes visible in neuronal somata within 2 days, reaching a maximum number of neurons and brightness within a week. We did not observe GFP expression in satellite cells in ganglia or Schwann cells along nerves. Soluble fluorescent proteins were transported along the axons of sensory neurons at a typical slow transport rate (2-3mm/day), reaching peripheral targets such as taste buds at the tip of the tongue in 2 weeks. We also confirmed that the PHP.S viral particles can be used for Cre-dependent expression for cell type selective targeting, and for functionally imaging select subtypes of sensory neurons.

Neuronal tracing of central circuits has emphasized the deliberate use of anterograde (from somatodendritic compartment down the axon) or retrograde (from axon terminal to soma) transported tracers. Over the last several decades, enzymes, dextrans, and viruses have been developed with directional selectivity. For AAV, particular natural and engineered serotypes have been shown to exhibit transport in one or the other, or both directions (McFarland et al., 2009; Aschauer et al., 2013; Rothermel et al., 2013; Castle et al., 2014). The directionality of AAV transport relies on microtubule-based mechanisms of directional transport in dendrites and axons (Leopold and Pfister, 2006). However, pseudo-unipolar sensory neurons such as those found in dorsal root, trigeminal, gustatory, and other visceral ganglia present a rather different version of polarity than that defined for central neurons (Nascimento et al., 2018; Shorey et al., 2021). These neurons have no dendrites, only a single axon that bifurcates with one axonal branch extending to a peripheral receptive field and the other towards central target(s). Thus, the directionality of AAV-PHP.S is difficult to define. With particles introduced into the circulation, the virus likely gains access directly to the somata of peripheral sensory neurons, based on our observation that reporter expression begins within 2 days of injection. The transport of GFP reporter seems to progress along both axonal directions at roughly similar velocity; GFP was detected along axons with progressively longer delay as distance from the soma increases (Fig. 4).

AAV-PHP.S was engineered with AAV9 as the parental capsid construct (Chan et al., 2017). AAV9 transduces astrocytes in adult brain but not enteric astrocyte-like glia (Gombash et al., 2014), and AAV-PHP.S was selected for expression in a GFAP-Cre mouse (Chan et al., 2017). Yet, we found that it did not transduce GFAP-expressing satellite glia in ganglia. We did observe some cells below lingual epithelium that were rapidly transduced (within 7 days of virus injection) that might be dendritic cells or Schwann cells (Fig. 4).

While many detailed connectivity maps of central pathways have been built using AAV, these tools have been applied much less to peripheral sensory neurons. Instead, mapping the spinal and brainstem targets of sensory ganglion neurons has relied on neuroanatomical tracing and well-defined molecular markers of neuronal types. This is the case even for recently identified sensory neuron subtypes (Haring et al., 2018; Wu et al., 2021). The use of AAV-PHP.S enhances the experimenter’s toolbox, allowing precise and efficient labeling of peripheral neurons while leaving resident neurons in the central target field unlabeled. With the recent molecular definition of peripheral gustatory neuron types, we anticipate this method will find substantial utility.

The use of Cre-dependent AAV-PHP.S constructs for targeting particular peripheral sensory neurons should open new directions for neuroanatomically and functionally defining sensory submodalities in the somatosensory as well as gustatory systems. For the taste system in particular, there is a critical need to define the brainstem targets of molecular classes of gustatory neurons. Controversies regarding the neuroanatomical basis for taste coding from the periphery to first central relays will benefit from these novel mapping tools. The resolution afforded by visualizing soluble, and synaptically targeted fluorescent reporters in individual fibers making synapses on defined peripheral receptors and rNST neurons (similar to Fig. 3E) will allow the development of accurate connectivity maps.

## Acknowlegments

This work was supported by NIH/NIDCD grants, R01 DC017303, R01 DC017303S1 and R01 DC018733 (N.C.); The Miami Project, The Buoniconti Fund and the State of Florida Red Light Camera Fund (P.T.). We thank Yania Ondaro-Martinez at the Miami Project Viral Core and Vivien Makhoul for help with video editing.

## FIGURE LEGENDS

**Extended Data Multimedia 1: Sensory fiber tracts in hindbrain.** 3-dimensional light sheet microscopy of hindbrain from mouse, 19 days after retroorbital sinus injection with AAV-PHP.S::CAG-mScarlet-I. Movie is a series of horizontal 2D images, representing a thickness of ≈1.5mm. AAV-delivered mScarlet fluorescence is enhanced with anti-RFP. Ascending spinal sensory afferents of the Gracilis and Cuneate tracts enter from the right. Cranial afferents readily visible include the spinal trigeminal tract (sp5) and gustatory afferents terminating in the nucleus of the solitary tract (NST).

**Extended Data Fig. 2:**
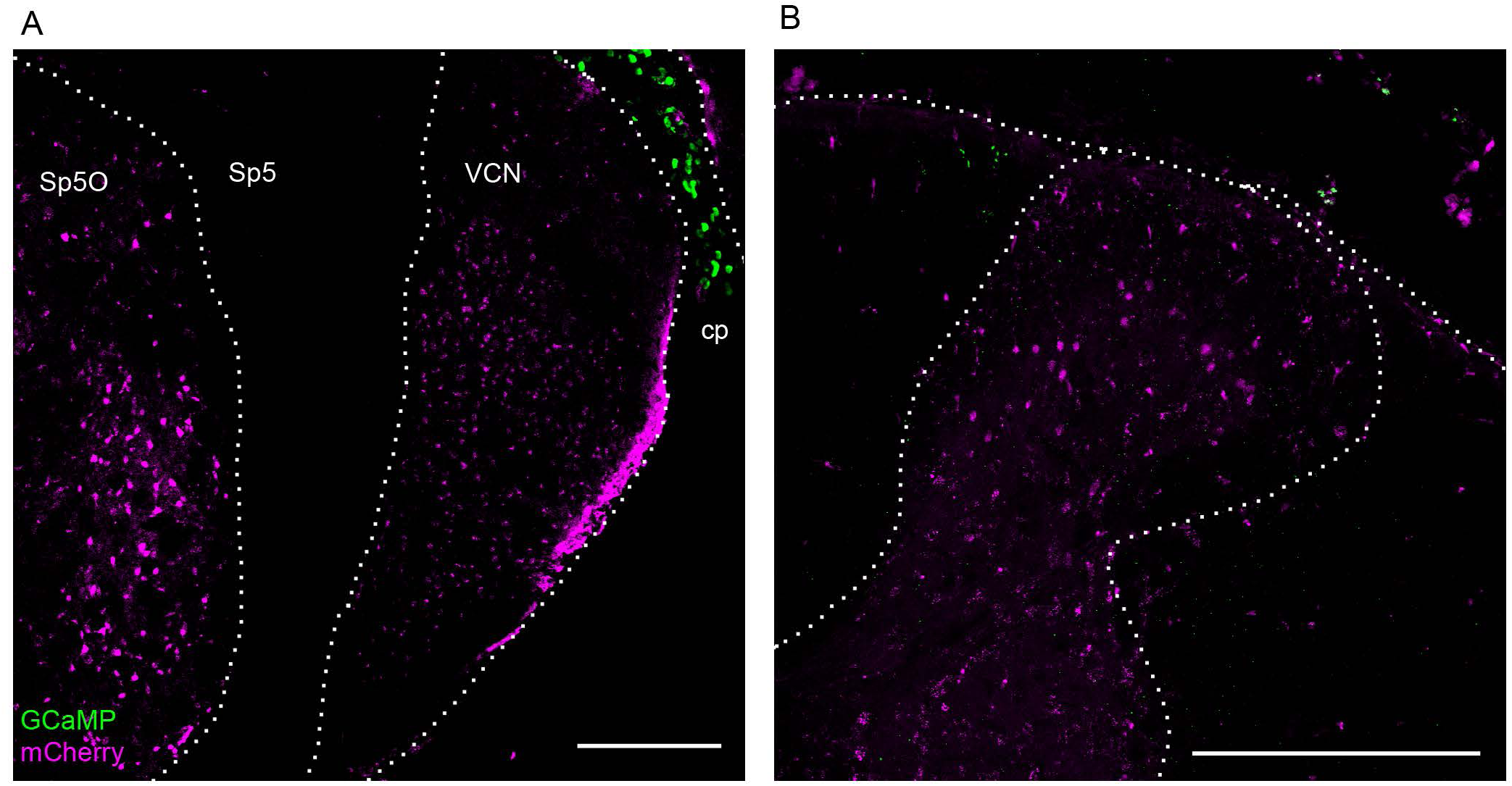
Cre-dependent expression is limited to peripheral neurons. a *Mafb-*mCherry-Cre mouse was injected with AAV-PHP.S::*flex*-GCaMP6s and perfusion fixed 19 days later. **A**, Cryosection of hindbrain shows mCherry+ (i.e. Cre-expressing) central neurons of the spinal trigeminal nucleus, oral part (Sp5O) and the ventral cochlear nucleus (VCN). These remain untransduced and completely devoid of GFP. Some GCaMP6 is visible in the choroid plexus (cp) within the lateral recess of the 4th ventricle. **B**, Cryosection of thoracic spinal cord similarly shows many Cre-expressing neurons of the spinal gray and these remain untransduced. In both cases, the peripheral afferents lack Mafb and Cre expression, and thus also are unlabeled. Scale bars, 250μm.

